# Design and Validation of an Open Loop Controlled Positive-Displacement Pump for Vascular Flow Modelling

**DOI:** 10.1101/2023.02.17.528991

**Authors:** Thomas O’Brien

## Abstract

Realistic vascular flow-rates are highly patient dependent. There is a need, when evaluating vascular medical devices, to study variable physiological flow-rates corresponding to specific patients. This study presents an experimental analysis of a positive-displacement piston pump designed for generating patient-specific vascular flow-rate wave-forms. Open loop control is used to drive a positive-displacement piston pump according to user-input physiological wave-forms. Transient and frequency analyses are performed for simple and complex inputs. Theoretical analysis of the system shows that it may be modelled as a linear first order system and the response of the system for various inputs shows that the real system approximates a linear first order response, however, higher order effects are noted. These higher order effects do not have any particular detrimental effect on the generation of physiologically realistic flow-rate waveforms.

## 1 Introduction

*In vitro* cardiovascular flow modelling requires versatile pumping systems capable of producing a variety of complex flow-rate waveforms. While versatile pumping systems provide researchers with the ability to model patient-dependant physiologically realistic flow-waveforms [**1**], limitations have been reported in the literature for certain pumping systems [**2**]. Numerous studies have used simple pumping systems to investigate cardiovascular flow but these approaches are unsuited toward patient-specific flows. Studies by Koslow *et al*. [**3**] and Viggers *et al*. [**4**] used peristaltic pumps capable of providing a certain flow-rate at a variable frequency. Other studies have used cam driven pumps [**5**], piston pumps [**6**] and centrifugal pumps [**11**]. Young & John [**7**] built and validated a versatile pump capable of producing variable physiologically realistic pressure pulses. In this study, a variable voltage input was used to produce a variable pressure output. However, the pump was not used to analyse variable flow-rates. A computer controlled gear pump was used to study blood flow characteristics in an *in vitro* phantom using power Doppler imaging [**12**]. A study by Liepsch *et al*. [**8**] used a complex pressure controlled system for producing variable flow-rate wave-forms, while Holdsworth *et al*. [**9**] used a computer controlled linear stepper to produce physiological flows. Stepper motors and microcontrollers have also been used in generating realistic flow conditions [**13**].

The difficulty associated with generating physiologically realistic waveforms is that the wave-forms are complex, represented by high-order polynomials or Fourier series. These higher order systems require programmable control systems to adequately represent patient-specific flows.

The objective of the study was to design and construct a programmable computer-controlled positive-displacement pump and to assess the resulting physiologically realistic flow-rate wave-forms. A theoretical model of the system was described and the transfer function for this model was derived. An associated control system to drive the pump was then developed. A stepper motor with high rotor inertia was used to drive a piston in contact with a working fluid according to a physiologically realistic pulse input by the user via a programming language. The transfer function for the theoretical model was then compared to that of the experimental model.

## 2 Theoretical Model

The theoretical model of the system is shown in fig. 1. A number of assumptions are made for this model, namely, (i) that the fluid is incompressible, (ii) no leakage occurs, (iii) the inertia of the piston and fluid are neglected, (iv) there are no viscous losses in the system, (v) the walls of the cylinder and pipes are rigid. A first order differential equation derived from the continuity equation describes the system as follows:

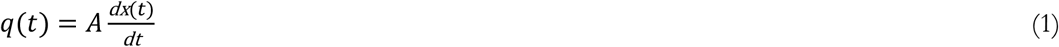

**Figure 1.**
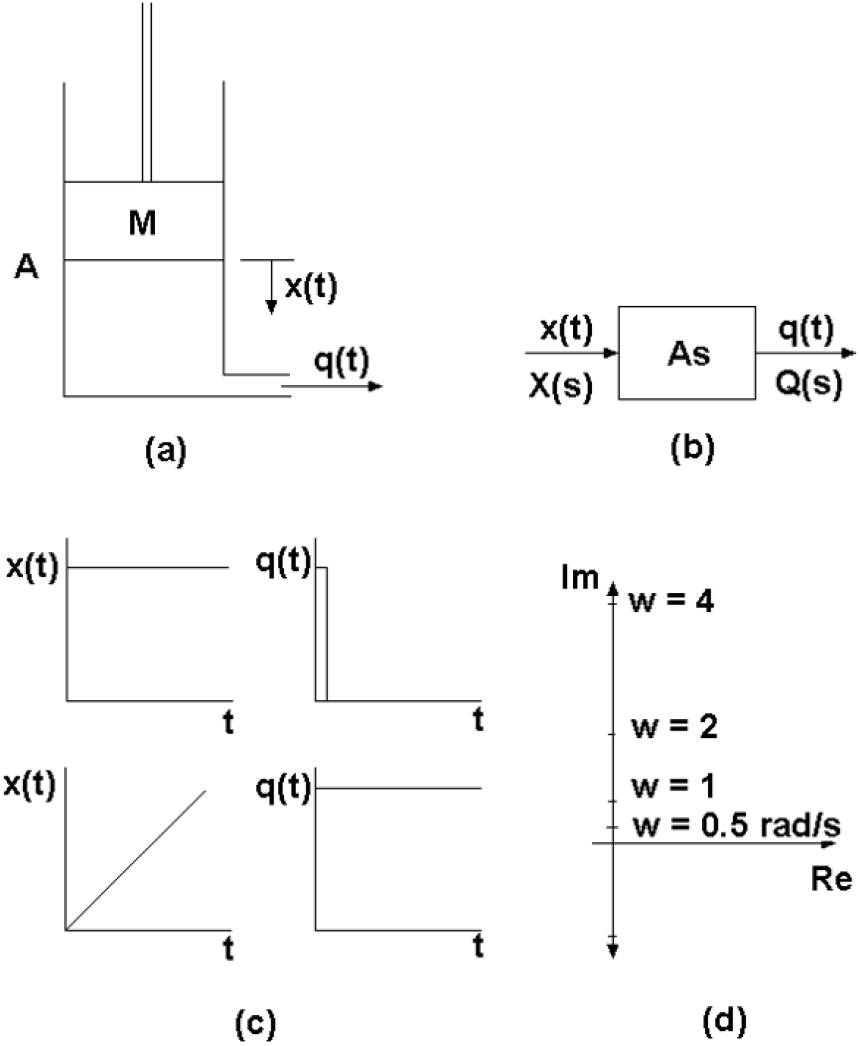
The theoretical model of the proposed system. (a) A position input, x(t), to the piston, M, results in an output flow-rate, q(t). (b) The open loop control and associated transfer function of the system. (c) The response of the system to step and ramp inputs. (d) The frequency response of the ideal system shows the output leading the input by 90°.

The response, q(t), of the system to a transient position input, x(t), is given by the following transfer function, fig. 1b:

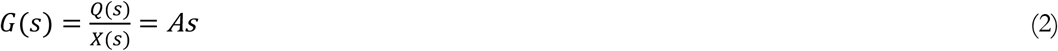

If the area, A, of the piston is unitary, it may be determined that the response of this system to a unit ramp input is a unit step output and the response of the system to a unit step input is a unit impulse, fig. 1c. The transfer function is that of a differentiator, therefore, the system differentiates input signals, fig. 1d.

## 3 Research Methodology

### Boundary conditions

The system was designed in order to produce a programmable pump that could reproduce waveforms occurring in major arteries such as the aorta. To achieve this, a physiologically realistic pulse [**10**] was used for the experimental test.

### Computer controlled piston pump

A positive displacement piston pump was designed and manufactured with a 30mm piston diameter (7.07×10^-4^m^2^) and a 96mm piston stroke length. One-way valves were mounted at the inlet and outlet of the piston chamber. The pump was mounted onto an aluminium base-plate and was coupled to the motor drive using an acetal resin rack-and-pinion with an 80mm pitch diameter.

A high rotor inertia (1400×10^-7^kg.m^2^) stepper motor (Vexta αStep ASM-98-AC ((Oriental Motor Europe)) was used to drive the piston. Motor resolution was 0.1 degrees/pulse. The stepper motor driver was connected to the computer using the DB 25 serial connector. Pins 2 (index counter-clockwise), 3 (index clockwise) and 4 (field excitation) were required to drive the motor with pin 25 used to ground. Pin 5 was used as a trigger for synchronisation with the flow-meter. A schematic diagram of the system is shown in fig. 2.

**Figure 2.**
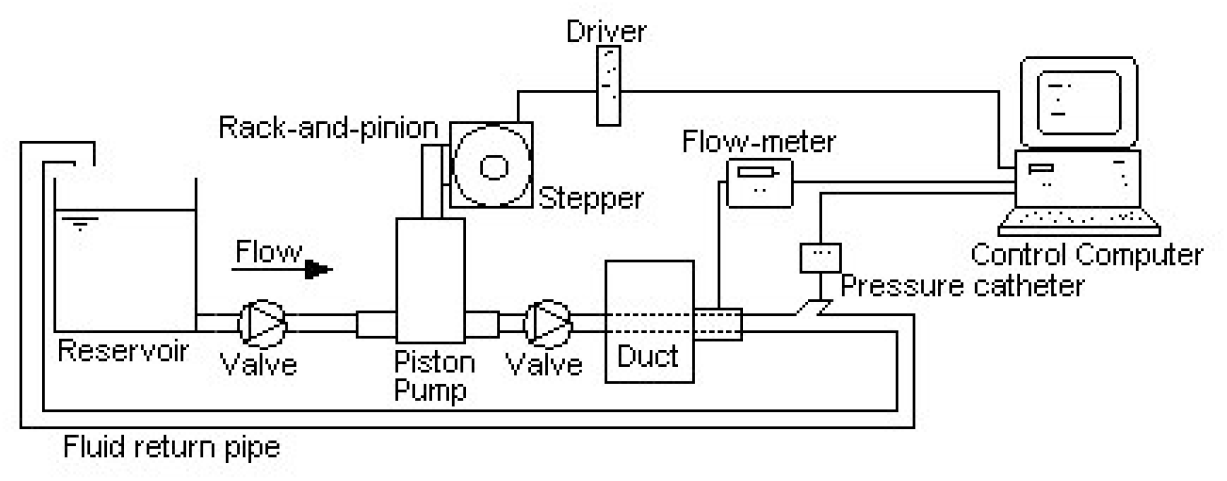
The programmable piston pump experimental set-up.

### Pulse train generation

A C++ program generates the digital pulse data necessary for the motor driver. Flow-rate wave-forms are converted into transient piston positional data which is then discretised to determine the required time-dependant angular positions of the motor. The effects of fluid inertia, friction, coupling backlash and stiffness are neglected and no signal preconditioning takes place. The algorithm used for the open loop control is shown in fig. 3. Briefly, a data file containing the time dependant flow-rate is read by the program. This flow-rate wave-form is then converted into time-dependant piston positional data. This data is discretised using a pulse timing delay factor and the piston position at each time interval is determined. Each digital pulse then indexes the stepper motor followed by the required delay. When the end of the pulse is found, the position plot is reversed and the piston returns to datum ready for the next stroke.

**Figure 3.**
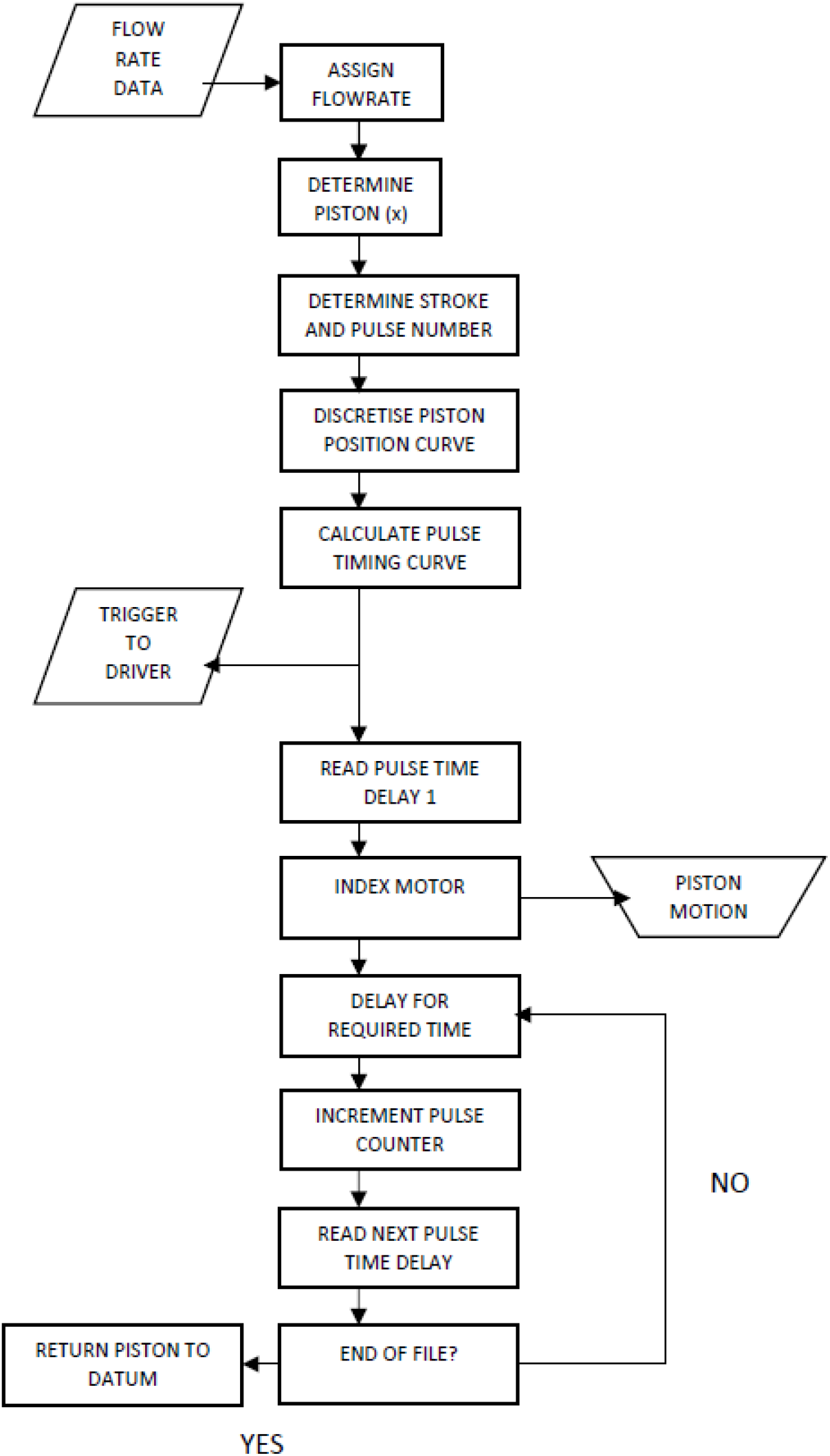
The computer algorithm used to control the piston pump.

### Validation flow circuit

The working fluid was water with an associated density of 1000kg/m^3^ and a viscosity of 0.01Pa-s. A 1m long transparent duct was used prior to entry to the flow-meter channel to enable the fluid to develop fully and also to act as a monitoring channel to detect the presence, if any, of air bubbles due to cavitation. The flow-meter was buffered from the piston-pump by a 1cm fluid contact length of compliant silicone rubber duct. This reduced vibration transmission through the walls of the system. An ultrasonic flow meter (Transonic Systems Inc.) was used to measure the flow-rate though a 20mm diameter pipe. The ultrasonic flow-meter system was calibrated to within 2% accuracy.

## 4 Results

The flow meter produces a linear output voltage signal which is calibrated for flow-rate measurements. The average flow-rate through the flowmeter was determined for each of the four ramp input frequencies. The frequency of 0.25Hz gave a flowrate of 1.77×10^-5^m^3^/s, 0.5Hz gave 3.58×10^-5^m^3^/s, 1Hz gave 7.04×10^-5^m^3^/s and 2Hz gave 13.4×10^-5^m^3^/s. The period of each waveform was also verified using the flowmeter.

### Response to simple one-stroke input

The response of the system to a ramp and a step input are shown in fig. 4. The ramp input is a steady 50mm piston stroke over 0.5s. The ramp input produces an approximate step output, though with significant oscillations. The maximum overshoot is approximately 54% with a rise-time of 0.035s (5%-95% step size). The steady-state flow-rate was 7.04×10^-5^m^3^/s. The overshoot suggested that higher order effects were present in the system.

**Figure 4.**
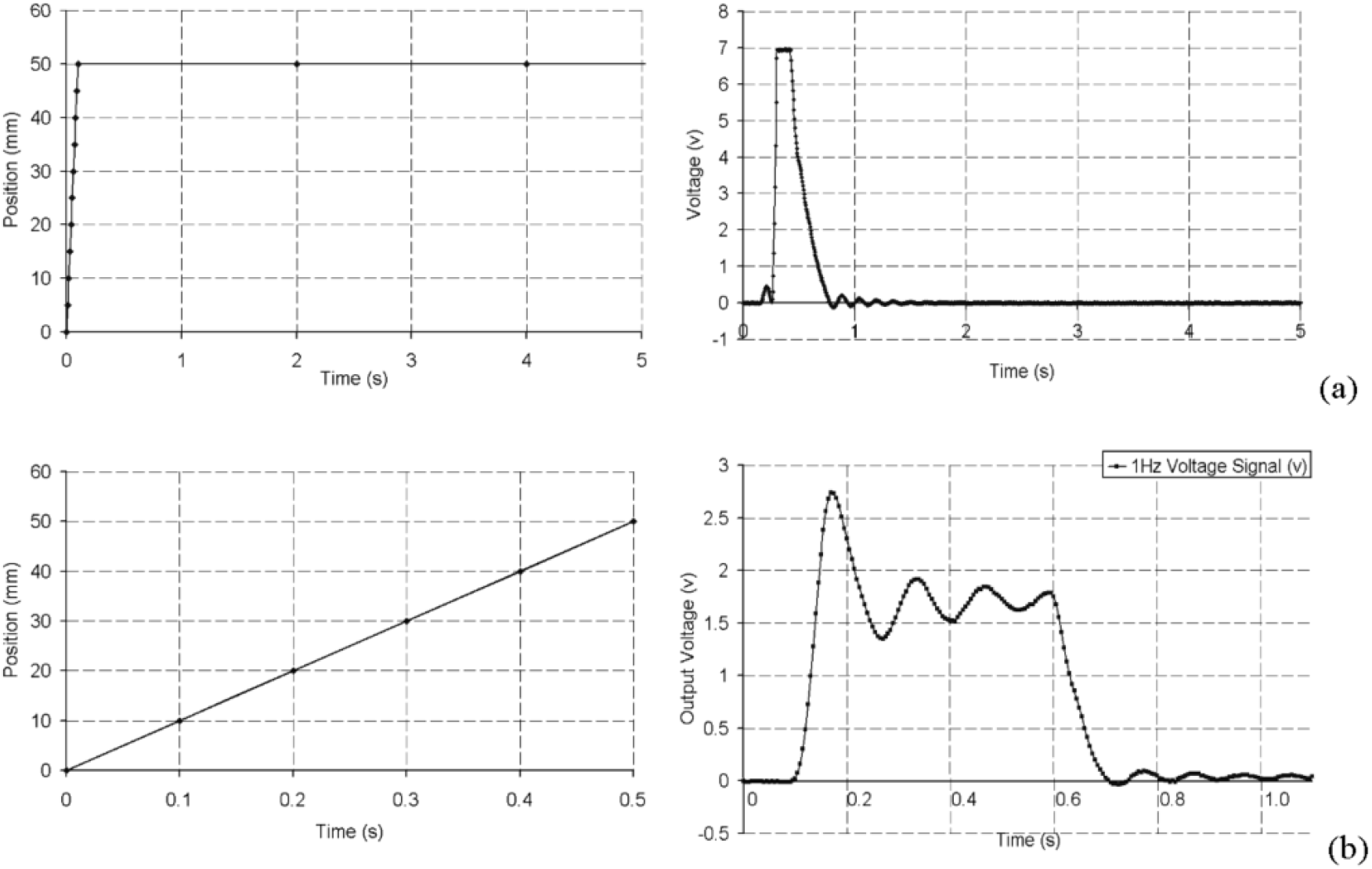
(a) A 50mm stroke over 100ms representing the impulse response test. The duration of the piston stroke was 100ms and the duration of the peak output pulse was 130ms. (b) Synchronised response of the system to a 1Hz ramp input. The steady stroke of 50mm is 0.5 seconds in duration and provokes an approximate step response by the system. There is a delay of approximately 0.1 second between the stroke of the piston and the measurement of the fluid response. Overshoot followed by three oscillations is evident.

An approximation of a step input was applied to the system. A piston stroke of 50mm was applied during a time interval of 100ms. A flow disturbance preceded the impulse output, likely caused by significant impact vibration of the piston due to rapid acceleration and deceleration. The rise-time of the impulse was approximately 0.05s. The peak flow-rate was calculated to be 28.3×10^-5^m^3^/s. The duration of the impulse maximum was 0.12s taking 0.35s to fall back to 0 volts with at least 6 oscillations following.

### Response to variable ramp frequency

This series of measurements involved recording the flow-meter output for four different ramp frequency inputs. For this test, the stroke length of the piston was maintained at 50mm and the input frequency was varied from 0.25Hz, 0.5Hz, 1Hz to 2Hz. The response of the system is shown in fig. 5. The peak maximum overshoots were 36%, 0.45s for the 2Hz signal, 34%, 0.45s for the 0.5Hz, 33%, 0.04s for the 0.25Hz and 54%, 0.35s for the 1Hz signal. The 0.25Hz pulse showed the presence of minor motor induced vibrations in the fluid and the ducts. While these were reduced, by compliant coupling of the duct to the piston pump, they were not eliminated.

**Figure 5.**
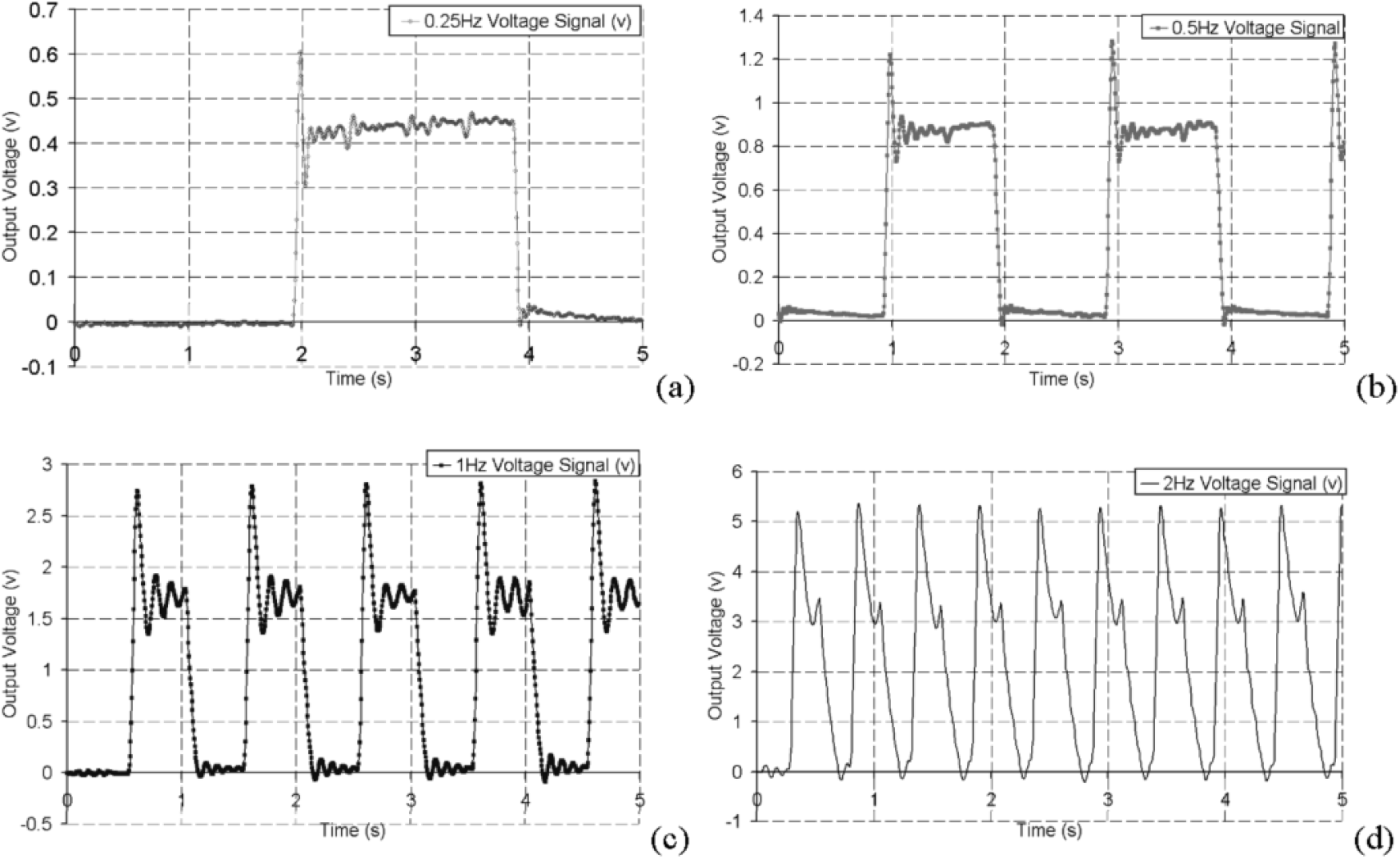
Samples of the measured output for a range of different input ramp frequencies. It may be seen that the 0.25Hz, 0.5Hz and 1Hz frequencies are approximately step responses with overshoot and oscillation though the signal becomes unsuitable at frequencies of 2Hz.

### Phase shift

A sinusoidal input of 1.07Hz with a 64mm stroke length, was applied to the system. The one-way flow valve at the piston outlet was removed and the valve at the inlet was sealed enabling oscillation of the fluid through the flow-meter. The position input was compared to the velocity output in fig. 6. The output sinusoid leads the input sinusoid by approximately 0.25s or 90 degrees phase. There are notable oscillations in the output signal. Integration of the output sinusoid yields a smoother pulse proportional to the displacement of the fluid in the flowmeter.

**Figure 6.**
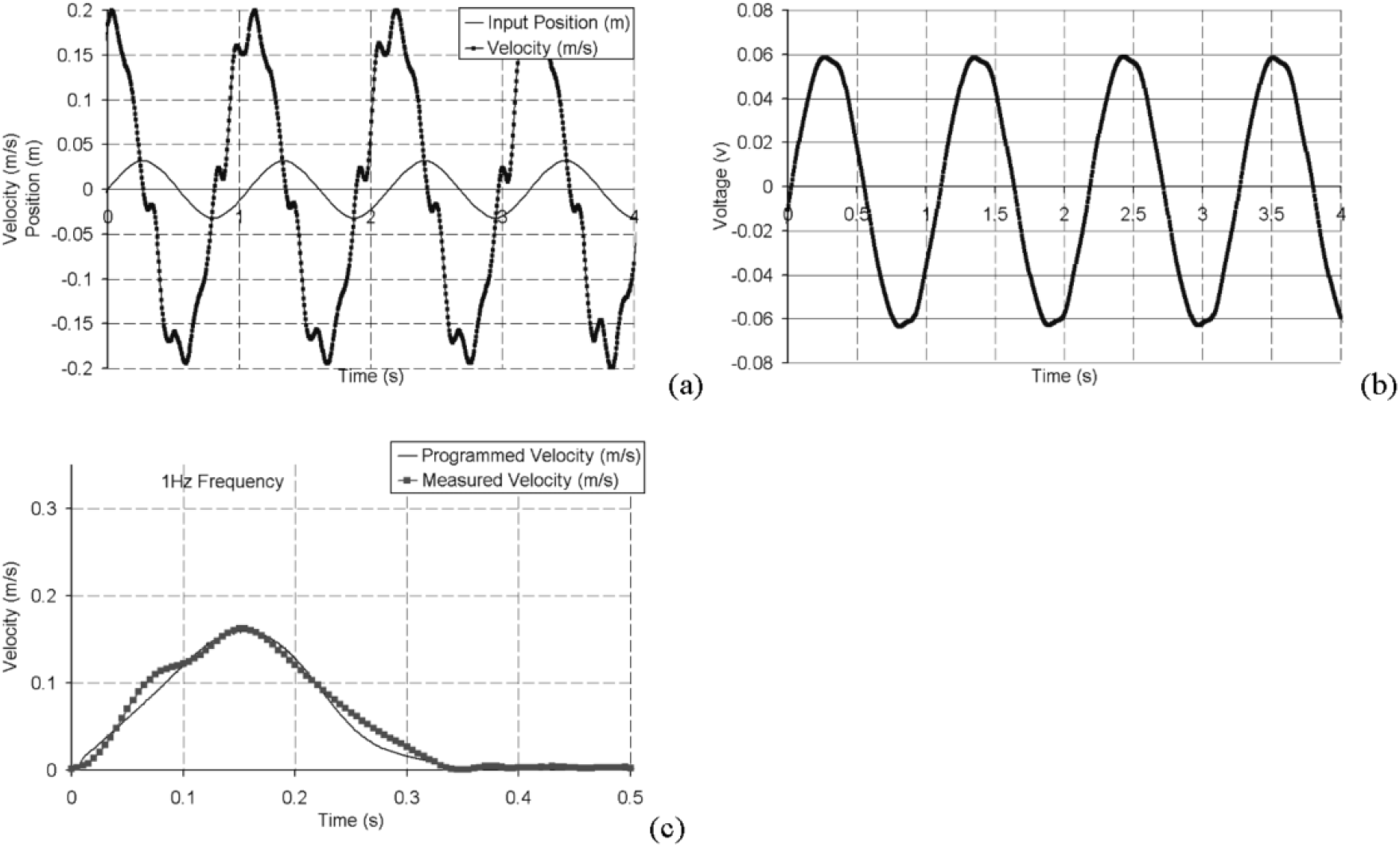
Response of the system to a sinusoidal position input (a). The steady state output velocity leads the input position by approximately 0.25s (90 degrees phase). There are noticeable oscillations in the output signal. Integration of the output shows a smoother signal (b). The response of the system to a physiologically realistic aortic velocity input for a frequency of 1Hz is shown (c). The 1Hz velocity output shows good agreement to the input though minor oscillations are evident.

### Physiologically Realistic Velocity Pulse

The system was used to model the physiologically realistic aorta flow-rate waveform [**10**]. The input velocity waveform and the resulting output velocity waveform compare favourably, fig. 6. It can be seen that the pumping system gives an excellent response to the 1Hz physiologically realistic pulse exhibiting only minor oscillations and an acceptable rise time.

## 5 Discussion

A simplified theoretical model was used to design a positive-displacement pumping system for reproducing physiologically realistic flow-rate waveforms. The transfer function of the simplified model was found to be a first-order linear differentiator. The associated transient and frequency responses were determined for simple inputs. The response of the real system showed acceptable approximation to the theoretical model for frequencies of up to 1Hz which corresponds to frequencies found in physiologically realistic resting flow-rate wave-forms, albeit with oscillations occurring. It is probable that these oscillations are caused by the neglect of secondary factors in the model assumptions. For example, the theoretical model neglected viscous drag forces due to the fluid. Modelling of viscous drag forces introduce a retarding factor in the governing equation:

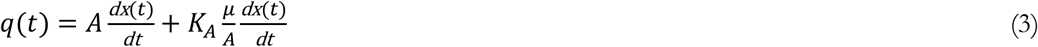

where the fluid pressure is, 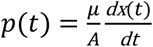 and the area coefficient is 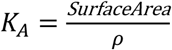.

Equation (3) illustrates how the differentiation rate is reduced by the presence of these viscous drag forces. In a similar manner, the assumption of incompressibility also removes an element from the transfer function. Compressibility is represented by:

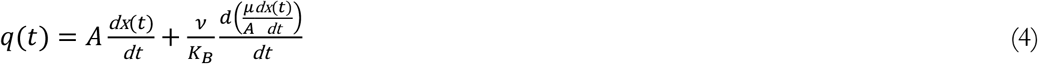

where ν is the kinematic viscosity of the fluid and *K_B_* is its bulk modulus.

This second order differential equation may be responsible for the introduction of second order oscillatory effects in the response of the real system.

Also neglected in the theoretical model were the effects of friction, valve oscillation, coupling flexibility, backlash and inertia. These may also lead to higher order effects which may explain the oscillations in the real system response.

In spite of the simplifications, the real system has shown to have an acceptable response to the applied inputs up to a frequency of 1Hz. Calculating the transfer functions at each of the four frequencies for the steady state outputs, shows acceptable approximation to the theoretical model, G(s) = 0.000707s. At 0.25Hz, G(s) = 0.000708s, at 0.5Hz, G(s) = 0.000716s, at 1Hz, G(s) = 0.000704s and at 2Hz, G(s) = 0.000670s.

The real system gives an acceptable response to the aortic flow-rate wave-form. The gradients of physiologically realistic waveforms for frequencies of up to 1Hz are not as severe as the ramp inputs applied for the theoretical model, and so, only minor oscillations are noted.

### Conclusion

An inexpensive open-loop control system is used to drive a positive-displacement piston pump and has been validated for physiologically realistic cardiovascular flows. The system is capable of producing variable flow-rate wave-forms as found in the cardiovascular system. The system approximates to a linear first order theoretical model for physiological frequencies of up to 1Hz. Higher order effects are noted in the response of the system, however, for physiologically realistic resting flow-rates, these higher order effects do not have a detrimental impact on the system response.

## Notes

### Competing Interest Statement

The authors have declared no competing interest.

## References

[1] M. Walsh, T. McGloughlin, D. W. Liepsch, T. O’Brien, L. Morris, A. R. Ansari, “On using experimentally estimated wall shear stresses to validate numerically predicted results”, Proc. Inst. Mech. Engrs., Journal of Engineering in Medicine, 217(H), 77–90, 2003. https://doi.org/10.1243/09544110360579286.

[2] M. M. de Jong, O. Parise, F. Matteucci, M. Rutten, M. Devos, M. Romano, L. R. Micali, G. Parise, J. G. Maessen, S. Gelsomino, “Aortic flow below and visceral circulation during aortic counterpulsation: Evaluation of an in vitro model”, Perfusion, Jan;37(1):69–77, 2022. doi: 10.1177/0267659120978641.

[3] A. R. Koslow, R. R. Stromberg, L. I. Friedman, R. J. Lutz, S. L. Hilbert, P. Schuster, “A flow system for the study of shear forces upon cultured endothelial cells”, Journal of Biomechanical Engineering, 108, 338–341, 1986. https://doi.org/10.1115/1.3138625.

[4] R. F. Viggers, A. R. Wechezak, L. R. Sauvage, “ ‘An apparatus to study the response of cultured endothelium to shear stress”, Journal of Biomechanical Engineering, 108, 332–337, 1986. https://doi.org/10.1115/1.3138624.

[5] Y. I. Cho, L. H. Back, D. W. Crawford, R. F. Cuffel, “Experimental study of pulsatile and steady flow through a smooth tube and an atherosclerotic coronary artery casting of man”, Journal of Biomechanics, 16(11), 933–946, 1983. https://doi.org/10.1016/0021-9290(83)90057-X.

[6] Z. D. Shi, S. H. Winoto, T. S. Lee, “Experimental investigation of pulsatile flow in tubes”, Journal of Biomechanical Engineering, 119, 213–216, 1997. https://doi.org/10.1115/1.2796082.

[7] S. T. Young & Y. C. John, “A flexible pressure pump for cardiovascular studies”, Medical Engineering and Physics, 17(4), 243–247, 1995. https://doi.org/10.1016/1350-4533(95)90848-6.

[8] D. Liepsch, G. Pflugbeil, T. Matsuo, B. Lesniak, “Flow visualisation and 1-and 3-D laser Doppler anemometry measurements in models of human carotid arteries”, Clinical Hemorheology and Microcirculation, 18, 1–30, 1998. Access online 16 February 2023 at https://pubmed.ncbi.nlm.nih.gov/9653582/

[9] D. W. Holdsworth, D. W. Rickey, M. Drangova, D. J. M. Miller, A. Fenster, “Computer controlled positive displacement pump for physiological flow simulation”, Medical & Biological Engineering & Computing, Nov, 565–570, 1991. Access online 16 February 2023 at https://link.springer.com/article/10.1007/BF02446086.

[10] C. P. Cheng, R. J. Herfkens, C. A. Taylor. “Abdominal aortic hemodynamics conditions in healthy subjects aged 50-70 at rest and during lower limb exercise: in vivo quantification using MRI”, Atherosclerosis, 168(2), 323–331, 2003. https://doi.org/10.1016/S0021-9150(03)00099-6.

[11] M. Schöps, S. H. Gros-Hardt, T. Schmitz-Rode, U. Steinseifer, D. Brodie, J. C. Clauser, C. Karagi-annidis, “Hemolysis at low blood flow rates: in-vitro and in-silico evaluation of a centrifugal blood pump”, Journal of Translational Medicine, 19, 2, 2021. https://doi.org/10.1186/s12967-020-02599-z.

[12] N. J. Raine-Fenning, N. M. Nordin, K. V. Ramnarine, B. K. Campbell, J. S. Clewes, A. Perkins, I. R. Johnson, “Determining the relationship between three-dimensional power Doppler data and true blood flow characteristics: an in-vitro flow phantom experiment”, Ultrasound Obstet Gynecol, Sep;32(4):540–50, 2008. doi: 10.1002/uog.6110.

[13] G. E. Engels, S. L. Blok, W. van Oeveren, “In vitro blood flow model with physiological wall shear stress for hemocompatibility testing-An example of coronary stent testing”, Biointerphases, Sep 18;11(3):031004, 2016. doi: 10.1116/1.4958979.

